# GenePy – a score for estimating gene pathogenicity in individuals using next-generation sequencing data

**DOI:** 10.1101/336701

**Authors:** E Mossotto, JJ Ashton, RJ Pengelly, RM Beattie, BD MacArthur, S Ennis

## Abstract

NGS is a revolutionising diagnosis and treatment of rare diseases. However, its relatively modest application in common diseases is limited by analytical approaches.

Instead of variant-level approaches, typical for rare disease or large cohort analyses, contemporary investigation of common polygenic disorders requires the development of tools combining mutational burden and biological impact of a personalised set of mutations into single gene scores. GenePy (https://github.com/UoS-HGIG/GenePy) is a gene score for transforming sequencing data capable of estimating whole-gene pathogenicity on a per-patient basis.

GenePy implements known deleteriousness metrics, incorporates allele frequency and individual zygosity information. Individuals harbouring multiple rare highly deleterious mutations accumulate extreme gene scores while the majority of genes usually achieve very low scores. Following correction for gene length, GenePy intuitively prioritises genes within individuals and affords gene/pathway score comparison between groups of individuals. Herein, we generate GenePy scores from whole-exome sequencing data for ∼15,000 genes across a cohort of 508 individuals. We describe score attributes and model behaviour under various biological conditions.

We demonstrate proof of concept that GenePy sensitively identifies known causal genes by calculating GenePy scores for *NOD2* (an established causal Crohn’s Disease gene), in a modest cohort of patients for comparison against controls. This test of GenePy using a positive control gene demonstrates markedly more significant results (p=1.37 × 10^-4^) compared to the most commonly applied tool for combining common and rare variation.

In addition to increasing the biological information content for each variant, the *per gene-per individual* nature of GenePy transforms the utility of sequencing data. GenePy scores are intuitive when assessing for individual patients or for comparing between groups. Because GenePy intrinsically reflects pathogenicity at the gene level, this specifically facilitates downstream data integration (e.g. into machine learning, network and topological analyses) with transcriptomic and proteomic data that also report at the gene level.

**Author Summary:** Rapid technological advances have made DNA sequencing an effective, economic tool for detecting genomic variation. Detecting rare variation at the individual level is proving very successful in identifying the genetic causes of disease when just single mutations are sufficient to manifest disease. However, interpreting genomic data is much less straightforward for common diseases such as asthma, arthritis or heart disease where many genetic changes across multiple genes combine with the environment to bring about disease symptoms.

We have developed a new scoring system called GenePy that generates whole gene pathogenicity scores for indiviual patients. The score corrects for the length of the gene and is intuitive to use. Unlike many mutation deleteriousness metrics, GenePy also takes into account the population frequency of the variant and the number of copies of any given mutation and combines data for as many variants as are present in a given gene for any one individual.

In this paper we apply the GenePy scoring system to a cohort of over 500 individuals for whom we have sequencing data across all genomics regions that code for protein. We descibe how GenePy performs and demonstrate superior sensitivity to detect known causal genes in a common autoimmune condition.

## Introduction

In the last decade, next-generation sequencing (NGS) has emerged as an effective tool for detecting single nucleotide variants (SNVs) causing rare conditions [1]. Recent retrospective studies have demonstrated an increase of 25-31% in diagnostic yield of rare diseases due to the application of exome or whole genome sequencing in a clinical framework [2, 3]. Through comparison against human genome reference sequence, high-quality NGS data on individual patients can be used to identify variation in variant call files (VCF). These files typically contain in excess of 30,000 variants when based on whole exome data that capture the coding region of the genome only and runs to many millions when based on whole genome data. The successful identification of disease causing variation is critically dependent upon annotation and subsequent filtering of these data. Filtering strategies typically focus on very rare variants in panels of genes empirically implicated as related to the clinical manifestation or phenotype of interest. Further exclusion of synonymous variants that have no impact on protein amino acid sequence and variants that occur at a frequency substantially greater than that of the disease of interest are also deprioritised. These steps can reduce the search space for causal variation by orders of magnitude to smaller sets of hundreds or even tens of genetic changes that are then prioritised by *in silico* methods[4].

Many *in silico* tools have been developed in order to estimate the potential impact of genetic variants on gene/protein function. Predicting pathogenicity or deleterious impact can be achieved through a variety of algorithms that focus on one or more specific biological aspect(s). Three broad classes of deleteriousness prediction metrics are: (i) conservation metrics, (ii) function alteration metrics and (iii) composite scores. Conservation metrics such as GERP++ [5], phastCons [6] and phyloP [7] assign a high deleteriousness to variants where the homologous position in other species has remained constrained over evolutionary history. Scores focused on predicting the potential disruption of protein functionality, for example through alteration of resultant protein amino acid sequence, include SIFT [8], FATHMM [9], fathmm-MKL [10], PolyPhen2 [11], MutationTaster [12], PROVEAN [13] and VEST3 [14].

To date, no single *in silico* metric has proven unilateral superiority in estimating consequent severity, despite an expanding list [15] of metrics based on subtly different foundations and assumptions. While individual metrics have the ability to perform well in isolation, discordant evidence when assessing the same data with multiple metrics has led to increased uncertainty in the choice of prediction tool [16]. This, in turn, has led to the development of a range of composite prediction tools applying statistical and machine learning methodologies that combine metrics assessing both conservation *and* functionality in order to obtain higher accuracy [17]. The most utilised composite scores include CADD [18], MetaSVM and MetaLR [19], M-CAP [20] and DANN [21] with no one method emerging as optimal [22]. For this reason, when assessing variant deleteriousness, it is still necessary to observe consensus prediction based on multiple scoring metrics rather than focusing on any single score [23]. This remains the case when studying rare Mendelian disease where single gene mutations imparting severe consequence are expected to represent the most extreme set of deleterious variants.

In contrast to rare diseases, common genetic diseases such as ischemic heart disease, asthma, inflammatory bowel disease (IBD) or Alzheimer’ disease are caused by the combined action of multiple genetic variants each differentially impacting risk and disease severity while working in combination with environmental exposures[24]. Collectively, common diseases impose an enormous economic burden and arguably have the greatest unmet need for diagnosis and stratified treatment [25]. The set of genes and variants imparting increased susceptibility vary from one patient to the next even when clinical presentation and molecular pathology appear indistinct.

Prior to transformative NGS approaches, genome-wide association studies (GWAS) made substantial advances in explaining the molecular bases of complex diseases. These studies tagged up to a million common single nucleotide markers across the genome and identified statistically significant distributions of biallelic markers in large cohorts of independent patients compared to ethnically match controls. Genetic regions implicated by GWAS were assumed to harbour genes or regulatory elements underpinning the disease of interest. However, because these genetic breakthroughs were achieved using necessarily huge cohorts of patients compared to controls, while their findings hold true for massive patient groups, they are largely uninformative on an individual patient basis. Importantly, the relevance and value of GWAS findings to individual patients has therefore not translated through to clinical practice in terms of either diagnosis or treatment.

Application of NGS to improve our understanding of common oligogenic diseases have been largely limited to burden tests that extend the association testing framework to integrate information about common and rare variation across discrete genomic regions such as genes. While this approach harnesses the power of NGS through the inclusion of rare variants that can only be detected by sequencing approaches, they are most often implemented through collapsing multiple variants into a single value for univariate analysis. The limited success of these approaches are partly attributed to their intrinsic lack of biological information and inclusion of both causal and benign genetic variation [26, 27]. In order to overcome this limitation, Neale *et al.* developed the C-alpha test, correcting for both protective and deleterious variants but at the cost of losing statistical power. Currently, SKAT (and SKAT-O optimised for small sample size) [28] represents the most sensitive approach to test for association between a genomic region and a phenotype. SKAT jointly assesses both rare and common variants maximising the statistical power and representing a new class of analysis lying between burden and association tests and has been successfully applied to a large variety of complex diseases [29–33].

While NGS is proving a transformative technology for the diagnosis and treatment of rare diseases, its relatively modest application in common diseases is limited by a lack of analytical approaches that incorporate *individual* profiles of genetic variation ascertained through NGS annotated with biologically meaningful information on their frequency and consequence.

Instead of variant focussed approaches typical for rare disease or large cohort approaches that distinguish GWAS, contemporary analyses of complex polygenic disorders requires the development of tools that combine both mutational burden and biological impact of a personalised set of mutations into single scores for discrete sub-genomic units such as genes. A matrix of such a set of scores for any one individual could then be analysed using various methodology including machine learning.

In this study, we describe the development and implementation of GenePy, a novel gene-level scoring system for integration and analysis of next-generation sequencing data on a per-individual basis. GenePy accounts for (any one of 16) variant pathogenicity scores, allele frequency and zygosity and sums across all variants within a gene for each patient. GenePy scores for a single gene can be compared between individuals. Following correction for gene size, all gene scores or subsets thereof can be implemented in downstream network analyses or used as input for machine learning to stratify disease subtypes. We validate GenePy sensitivity by comparing the genes scores for a cohort of paediatric IBD patients against a non-IBD cohort for the *NOD2* gene – a widely accepted positive control gene for causality in complex IBD.

## Methods

### Sample data

Whole exome sequencing (WES) data were derived from two sources. This first group comprised 309 patients diagnosed in childhood with IBD. This cohort (further described in [34]) includes unrelated, Caucasian patients ascertained and recruited through Southampton Children’s Hospital who were diagnosed under the age of 18 years according to the modified Porto criteria [35]. Additional WES data from a cohort of 199 anonymised individuals diagnosed with an infectious disease but unselected for any form of autoimmune disease were also used to give a total cohort size of 508 individuals with WES data.

Genomic DNA was extracted from peripheral venous blood and fragmented DNA subjected to adaptor ligation and exome library enrichment using the Agilent SureSelect All Exon capture kit versions 4, 5 and 6. Enriched libraries were sequenced on Illumina HiSeq systems.

### WES data processing

Raw sequencing fastq sequencing data from all 508 samples were processed using the same custom pipeline. VerifyBamID [36] was utilised to check the presence of DNA contamination across our cohort of 508 individuals. Alignment was performed against the human reference genome (GRCh38/hg38 Dec. 2013 assembly) using BWA [37] (version 0.7.12). Aligned BAM files were sorted and duplicate reads were marked using Picard Tools (version 1.97). Following GATK v3.7 [38] best practice recommendations [39], base qualities were recalibrated in order to correct for systematic errors produced during sequencing. Finally, variants were called using GATK HaplotypeCaller was applied to produce a gVCF file for each sample. Samples were processed on the University of Southampton IRIDIS cluster requiring an average of 4 hours run time per sample on a 16-processor node.

While the standard VCF format reports only alternative calls, the gVCF format identifies non-variant blocks of sequencing data and returns reference calls for loci therein. This enables affirmative calling of homozygous reference loci when combining call sets from multiple samples. Multi-sample variant calling was achieved through calling each individual sample separately and then merging all gVCFs using GATK GenotypeGVCFs. Processing efficiency was optimised for the set of 508 individual samples through batching into six subsets using GATK’s CombineGVCFs (approx. 6 hours/batch on a 16 processor node) and the resultant six gVCF files were merged for genotyping with GenotypeGVCFs (approx 1h on a 16 proc. node). Annotation of this composite file applied Annovar v2016Feb01 using default databases refSeq gene transcripts (refGene), deleteriousness scores databases (dbnsfp33a) and dbSNP147). Variant allele frequencies were sourced through Annovar (ExAc03[40]) or ensembl human variation API [41] where ExAc data were missing.

### Quality Control framework

In order to reduce heterogeneity, it is necessary to control for bias encountered due to alternative capture kit versions and variant quality. For the entire cohort of 508 samples, exon enrichment was performed using Agilent SureSelect capture kits but at different time-points. For this reason, there is inter-capture kit variability across the 508 cohort with kit versions 4, 5 and 6 being applied. To correct for disparity in the regions targeted by respective versions, all downstream analyses were restricted to the set of overlapping targeted genomic locations (as defined by respective kit BED files) using BEDtools v2.17 [42].

Following GATK best practice guidelines, HaplotypeCaller default settings were utilised, implying that only variants with a minimum Phred base quality score of 20 were called.

### GenePy Score

Individuals typically have multiple variants across the coding region of genes making the interpretation of their combined effect challenging. We hypothesised that for each individual sample *h* within our cohort *H = {h*_*1*_, *h*_*2*_, *…, h*_*n*_*},* the loss of integrity of any given gene *g* in the refGene database *G* = {g_1_, g_2_, … g_m_} can be quantified as the sum of the effect of all (*k*) variants within its coding region observed in that sample, where each biallelic mutated locus (*i*) in a gene is weighted according to its predicted allele deleteriousness (*D*_*i*_), zygosity and allelic frequency (*f*_*i*_). The GenePy score *S*_*gh*_ for a given gene (*g*) in individual (*h*) is

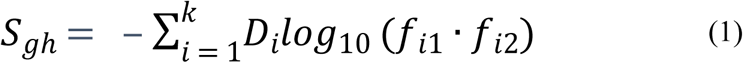

Importantly, the choice of variant deleteriousness score is user-defined, and therefore the GenePy score is able to take into account different definitions of pathogenicity depending on context. Herein we examine the relative attributes of using any one of sixteen of the most commonly applied scores (Table 1). At any one variant locus (*i*), we represent both parental alleles using *f*_*i*1_ and *f*_*i*2_ to embed the population frequency of allele_1_ and allele_2_ and in doing so model observed biological information on both frequency and zygosity. Any homozygous genotype therefore is simply the observed allele frequency squared whereas the product of each of the observed alleles is calculated for heterozygous genotypes. The latter can therefore accommodate variant sites with multiple alleles in addition to the typically encountered bialleleic single nucleotide polymorphisms (SNPs). Hemizygotic variation from male X-chromosomes are treated as homozygotic. Where a variant may be novel to an individual or absent from reference databases, we impose a lower frequency limit of 0.00001. This lower limit is arbitrarily set to conservatively reflect the lowest frequency that can be observed in the largest current repositories of human variation (ExAc03). The log function is applied to upweight the biological importance of rare variation.

**Table 1.**
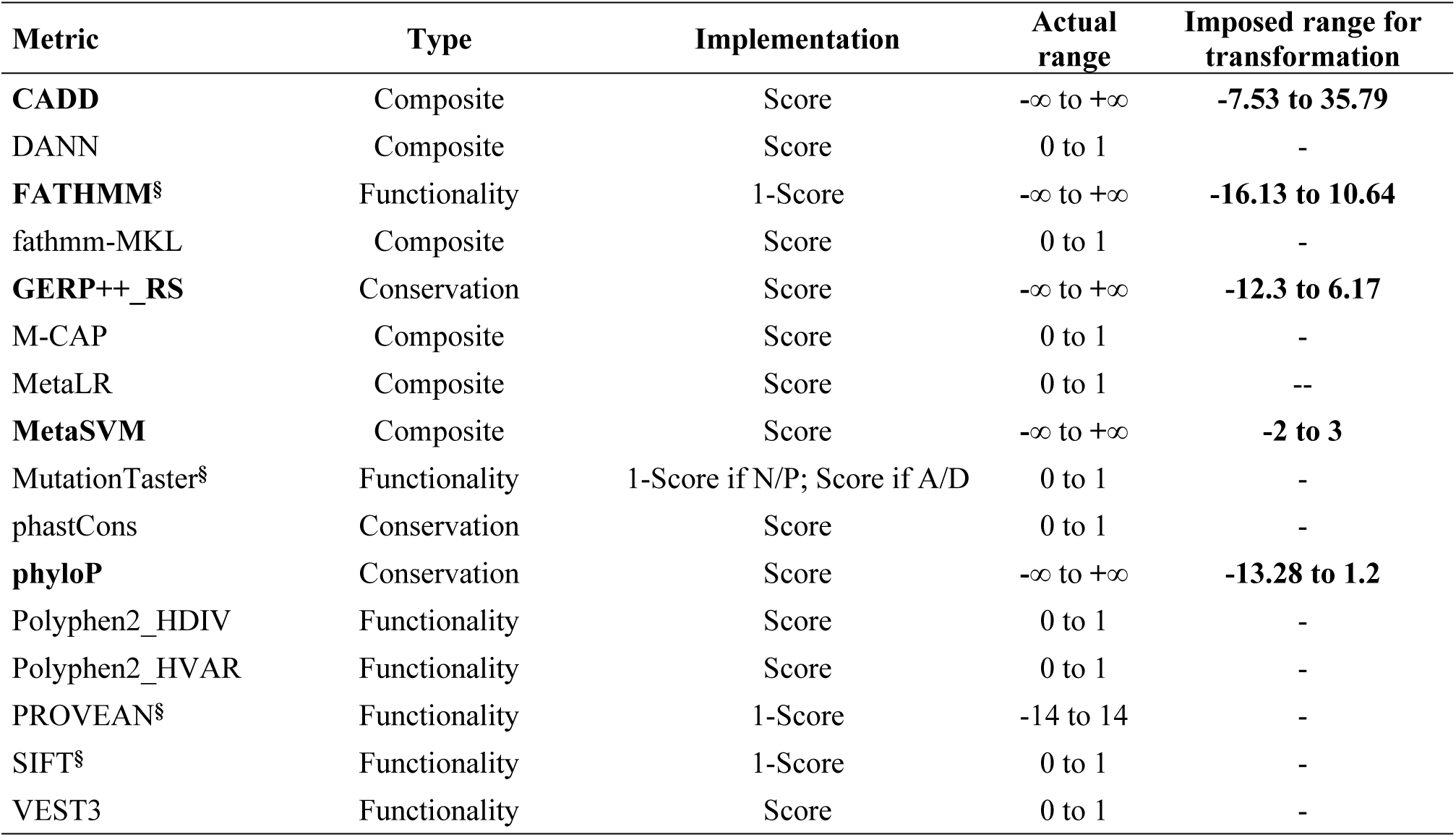
Pathogenicity scores for SNVs and their reported ranges in the dbsnfp database. §In order to maintain uniform directionality, the complement (1 – score) of a value was taken so that across scores, a value of 0 consistently indicated benign variation and a value of 1 inferred maximal pathogenicity.

| Metric | Type | Implementation | Actual range | Imposed range for transformation |
| --- | --- | --- | --- | --- |
| **CADD** | Composite | Score | $-\infty$ to $+\infty$ | **-7.53 to 35.79** |
| DANN | Composite | Score | 0 to 1 | - |
| **FATHMM<sup>§</sup>** | Functionality | 1-Score | $-\infty$ to $+\infty$ | **-16.13 to 10.64** |
| fathmm-MKL | Composite | Score | 0 to 1 | - |
| **GERP++_RS** | Conservation | Score | $-\infty$ to $+\infty$ | **-12.3 to 6.17** |
| M-CAP | Composite | Score | 0 to 1 | - |
| MetaLR | Composite | Score | 0 to 1 | -- |
| **MetaSVM** | Composite | Score | $-\infty$ to $+\infty$ | **-2 to 3** |
| MutationTaster<sup>§</sup> | Functionality | 1-Score if N/P; Score if A/D | 0 to 1 | - |
| phastCons | Conservation | Score | 0 to 1 | - |
| **phyloP** | Conservation | Score | $-\infty$ to $+\infty$ | **-13.28 to 1.2** |
| Polyphen2_HDIV | Functionality | Score | 0 to 1 | - |
| Polyphen2_HVAR | Functionality | Score | 0 to 1 | - |
| PROVEAN<sup>§</sup> | Functionality | 1-Score | -14 to 14 | - |
| SIFT<sup>§</sup> | Functionality | 1-Score | 0 to 1 | - |
| VEST3 | Functionality | Score | 0 to 1 | - |

Deleteriousness metrics were developed to assess damage induced by nonsynonymous variation, therefore structural variants such as frameshifts or stop mutations that truncate proteins are not routinely assigned deleteriousness values. Due to their highly detrimental impact to function we assign all protein truncating mutations the maximal deleteriousness value of 1. Synonymous and splicing variants are not routinely annotated by ANNOVAR and were not included in the current assessment.

Sixteen of the most common deleteriousness (D) metrics were selected for implementation within the GenePy algorithm (Table 1). Five of these metrics (shown in bold) are unbounded. In order to implement unbounded metrics in GenePy it was necessary to impose lower and upper limits by applying the respective minimum and maximum values observed in the dbnsfp33a database of 83,422,341 known SNV mutations. These limits were used to transform observed values in our cohort scaled to 0-1.

As a function of their size alone, larger genes have greater opportunity to accrue higher deleterious GenePy scores through having a greater number of variants thus inflating GenePy scores. We therefore generated GenePy scores corrected for the length of targeted gene regions (GenePy_cgl_) by dividing the GenePy score by the targeted length in base pairs and then multiplying by the median observed targeted gene length in our data (1461 base pairs). A final set of 16 deleteriousness metrics, each with a range of 0-1 where highest values were most deleterious, were individually implemented in the model.

### Score validation

In the absence of any comparable gene-based scoring system for individuals, GenePy performance was benchmarked by assessing its power to determine significantly different score distributions in disease cases compared to controls for a known causal gene and using the same variant data, comparing GenePy results against that of SKAT-O - the most commonly applied gene-level association test. The cohort comprised 309 individuals diagnosed with inflammatory bowel disease (IBD) and 199 controls unselected for autoimmune conditions. The analysis focussed on the *NOD2* gene - the most strongly and repeatedly associated common disease gene conferring strong association specifically with the Crohn’s disease (CD) subtype of IBD [43–45]. *NOD2* was selected as a positive control gene, whereby evidence for increased burden of deleterious mutation encoded in CD patient DNA compared to either ulcerative colitis (UC) or control DNA is expected.

The matrix of *NOD2* GenePy scores calculated for all 508 samples was split into controls and cases with the latter further divided into UC and CD subtypes. Statistical significance of GenePy score distribution difference between groups was calculated using the Mann Whitney U test for unpaired data. Using the same variant input data, the SKAT-O gene-based test for association was performed twice using default settings: firstly by considering all variants called within *NOD2* and secondly including only rare variants (MAF<0.05) as per developer recommendations [28].

Association tests succumb to false positive results due to spurious association brought about by population stratification or systematic differences in case versus control data. We excluded non-Caucasian individuals identified through comparison against the 1000 Genomes Project [46] using Peddy software [47] for ethnic imputation. We enforced parity in sequencing depth (known to impact power to call genetic variation [48]) for case-control data by limiting all score validation data to variants called in gene regions with a minimum read depth of 50X.

## Results

### QC results

All WES data (n = 508, n_ibd_ = 309, n_ctrl_ = 199) underwent quality control assessment for contamination using VerifyBamID and were confirmed free of contamination (free-mix statistic < 0.01). Out of 508 individuals, we identified three pairs of first degree relatives, one set of monozygotic twins and one mother-father-child trio. In order to correct for relatedness, which would bias association tests, for each pair, the sample with poorest coverage data was excluded. For the trio, the child data were excluded and unrelated parents retained.

### GenePy score behaviour – impact of allele frequency and zygosity

Figure 1 shows the results of simulated GenePy score (y-axis) calculated across a range of deleterious metric scores (0.1, 0.5, 0.75, 0.9, 0.95, 0.99) with varying minor allele frequency (x-axis) and further depicts the consequence of heterozygote versus homozygote states. The plot reveals the logarithmic nature of GenePy scores for a single locus only (whereas for any individual, their per gene GenePy score is weighted sum of all variant scores observed in that individual across that gene). For any single variant, the theoretical maximum observable GenePy value of ten occurs only with highest deleteriousness value (*D*), the lowest minor allele frequency (MAF = 0.00001) and in the homozygous state whereas the upper limit for a heterozygote with the same deleteriousness and frequency settings is five. The logarithmic scale implemented in GenePy algorithm confers rapidly increasing scores as the MAF approaches novelty.

**Figure 1.**
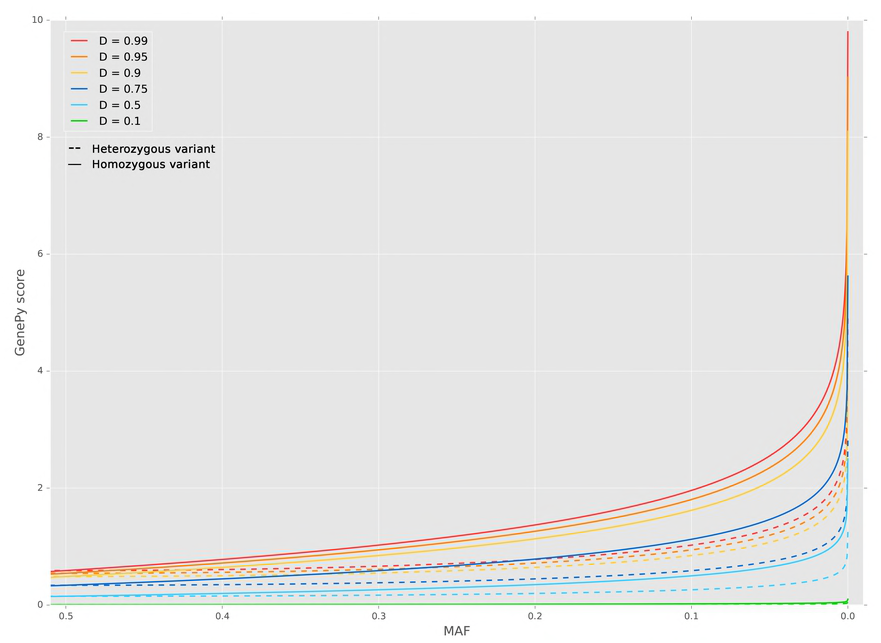
Single variant GenePy score distribution under fixed deleteriousness values. Impact of varying zygosity and minor allele frequency (MAF).

### GenePy score behaviour – impact of deleteriousness metric

While there are 27,238 genes annotated in RefSeq, we aimed to generate GenePy scores only for the overlapping subset of 21,577 target genes captured by all versions of the SureSelect capture kits applied. The GenePy scoring algorithm was executed for each of sixteen commonly applied metrics (Table 1). There is fluctuation in the number of genes for which variants were annotated with deleteriousness metric data using ANNOVAR ranging from 12,921 for M-CAP (one of the most recently released scores) to 14,745 genes annotated scores for Polyphen2_HDIV (one of the earliest developed deleteriousness scores) (Table 2). Among the 508 individuals that underwent GenePy scoring of exome data, the majority of genes are invariant within any one individual (e.g. median 9917 for CADD metric). This is expected for intrinsically sparse genomic data. However, across the cohort, no single gene returns a GenePy score of zero in all individuals indicating all genes have at least one rare variant observed amongst the 508 individuals. The vast majority of genes are scored with GenePy values of less than 0.01 and correction for gene length marginally increases the number of genes achieving lowest scores. More than 97%of genes achieve a score of less than 0.01 when the M-CAP metric is used whereas FATHMM scores approximately 65%of genes in the 0 –0.01 range. The inflated percentage of invariant genes observed when implementing M-CAP is explained by its tendency to depress weight for benign variants compared to other tested metrics [20].

**Table 2.**
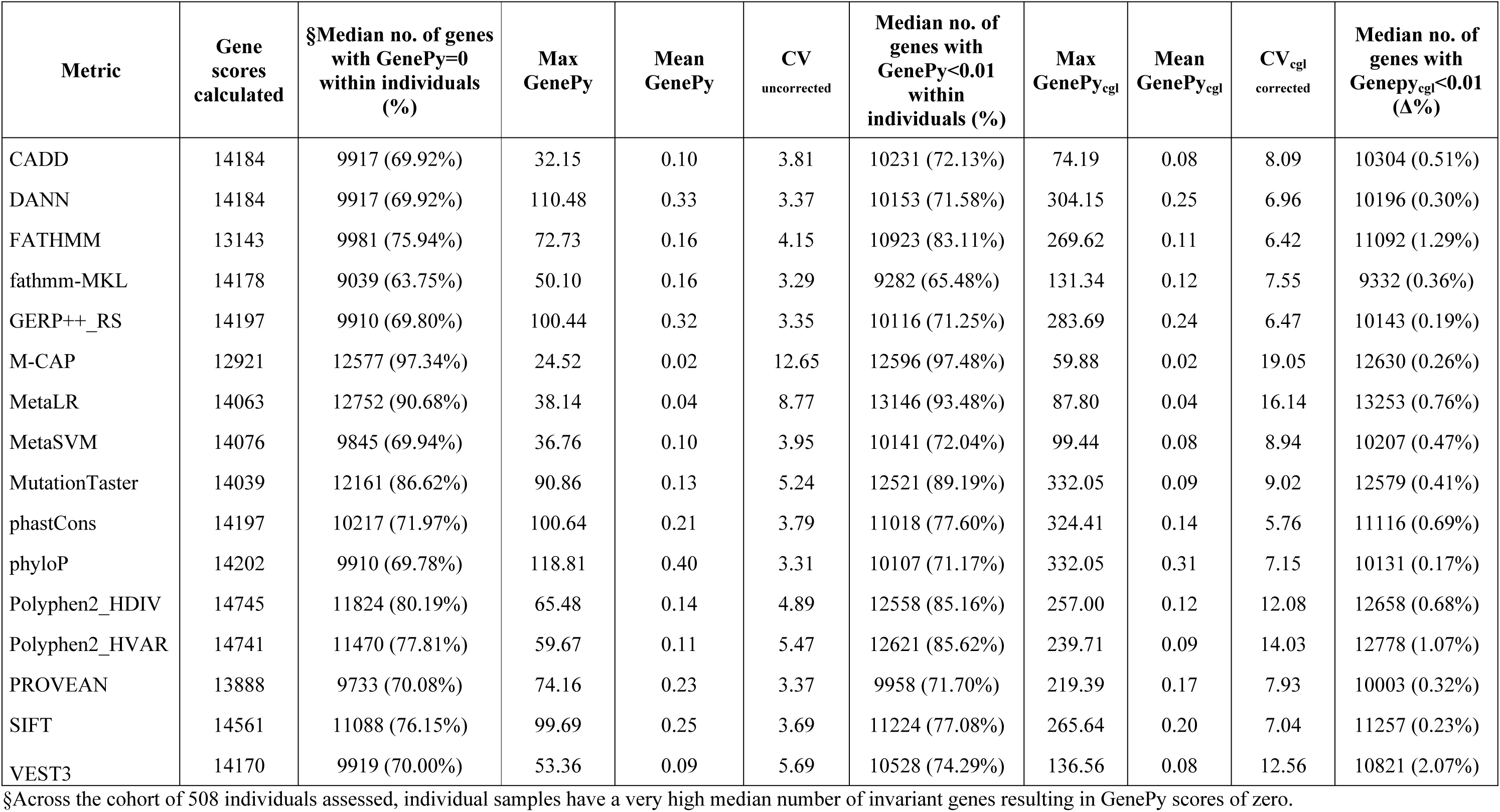
Statistical attributes of whole gene GenePy scores computed for sixteen deleteriousness metrics. Number of genes for which GenePy scores were calculated, median number of non-variant genes (GenePy=0), mean GenePy scores, mean and standard deviation across our cohort (n=508), coefficient of variation (CV, defined as σ/μ) and the median number of genes with a GenePy score <0,01 as percentage of the total number of genes. The same information is reported for GenePy_cgl_.

Across the ∼14,000 genes achieving GenePy scores, the observed score mean (uncorrected for length) in our cohort of 508 samples ranges from 0.02 to 0.40 depending on the applied deleteriousness metric. There is only modest effect on the range of the mean scores observed after correction for gene length (0.02 – 0.31). However, the gene length correction causes increased spread of the data reflected by an approximate two-fold increase in the coefficient of variation (CV) for GenePy scores generated that is consistent across all sixteen deleteriousness metrics. GenePy scores generated with M-CAP are least impacted by gene length correction but maintain the largest CV despite this score demonstrating the lowest maximum value.

In order to further investigate the behaviour of GenePy scores across genes, we calculated the median number of genes exhibiting scores falling within non-overlapping bins across the entire cohort. Figure 2 shows the profiles for the 0.01 to 6 range of GenePy scores and a bin size of 0.01. Genes with scores <0.01 are overrepresented (Table 2) and not shown. Across most of the sixteen metrics, a distinct pattern characterised by two spikes around uncorrected GenePy scores of 0.6 and 5 represent genes strongly influenced by a single highly deleterious common homozygous variants (*D=1*, MAF=0.5) or a single highly deleterious very rare heterozygous variant (*D=1*, MAF=0.00001) respectively. This profile was apparent for most deleteriousness metrics (except CADD, FATHMM, MetaSVM and VEST3, see Supplementary Figure 1). These two distinctive spikes are not observable once GenePy scores are corrected for the targeted gene length (Figure 1, lower panel and Supplementary Figure 2). We did not observe further spikes or other anomalies in the long right tail of the distribution of scores greater than 6.

**Figure 2.**
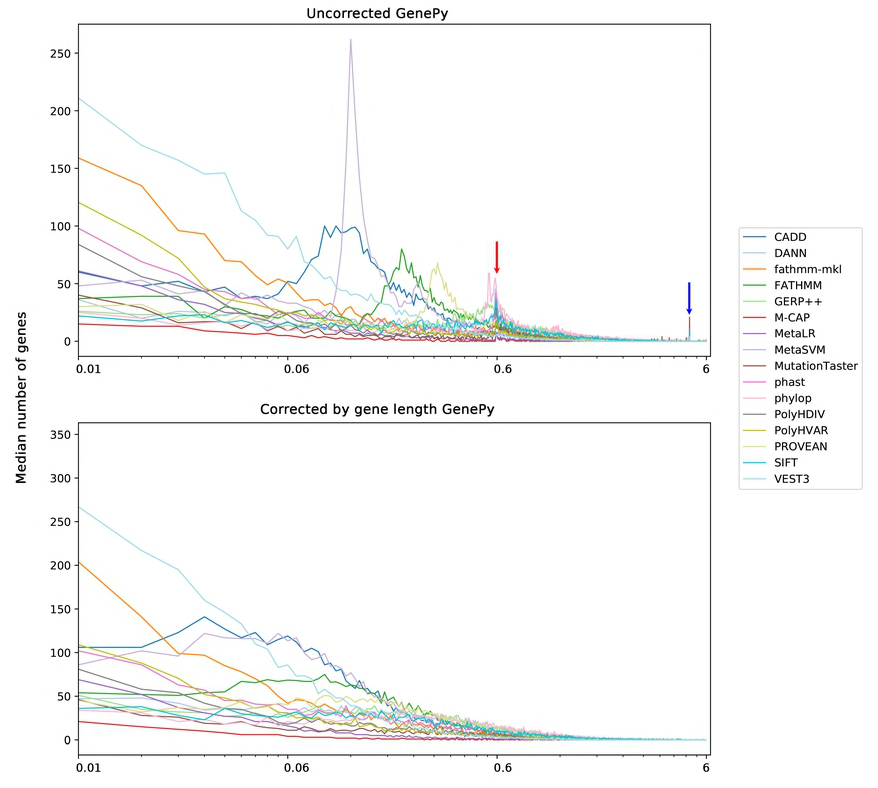
GenePy profiles observed for all genes across the whole cohort for all sixteen deleteriousness metrics. Uncorrected GenePy scores (upper panel) exhibit characteristic spikes reflecting gene scores strongly influenced by the effect of: single highly deleterious (D = 1) common homozygous variants (red) or; single highly deleterious very rare/novel variants (MAF = 0.00001) (blue). GenePy_cgl_ score profiles (lower panel) do not display these spikes. Invariant genes conferring a GenePy score <0.01 are overrepresented and not shown here by commencing the x-axis with the 0.01-0.02 bin. All sixteen versions of the GenePy score exhibit long tails in the GenePy score distribution truncated here at a score of six.

### GenePy score testing

Bias conferred by *NOD2* gene coverage, related samples and non-Caucasian ethnicity (Supplementary Figure 3) was removed from all IBD cases (n = 6_<50x_, n = 1_relative_ and n =20_non-Caucasian_) and non-IBD control samples (n = 16_<50x_, n = 4_relatives_ and n =13_non-Caucasian_) respectively. There remained 282 IBD cases for analysis of which 172 were diagnosed with Crohn’s disease, 100 with ulcerative colitis and a further 10 patients had a diagnosis of IBD undetermined (IBDU). There was a corresponding number of 166 controls.

The *NOD2* GenePy scores for the 282 IBD and 166 control individuals were calculated using all sixteen deleteriousness metrics. (Supplementary Figure 4). Given *NOD2* gene variant association is specific to the CD subtype of IBD, we calculated GenePy scores for both subtypes and grouped separately (Supplementary Table 1).

The Mann-Whitney U test comparison of the distribution of *NOD2* GenePy scores between all IBD, CD and UC subtypes against controls identified statistically significant differences (Table 3). Only modestly significant differences for just three of the implemented deleteriousness metrics (M-CAP, fathmm-mkl and MutTaster) were observed comparing all IBD against controls in this relatively small sample. When the cases were stratified by disease subtype, UC samples had significantly lower GenePy scores compared to controls but only for two of the implemented deleteriousness metrics (MetaLR, phastCons). As expected, the most significant difference in *NOD2* score distribution was observed when comparing CD patients only against controls. Without exception, a highly significant difference was observed using every deleteriousness metric with M-CAP the most significant (p = 1.37 × 10^-4^) all of which would withstand correction for the three independent tests performed. Regardless of which deleteriousness metric is used, the mean GenePy score is consistently higher in CD patient when compared with controls.

**Table 3.** NOD2 GenePy score statistics (maxima and means) and Mann-Whitney U tests across groups for all sixteen deleteriousness metrics. p-values smaller than 1 × 10^-2^ or smaller than 5 × 10^-2^ are highlighted by two (**) or one (*) asterisks respectively. SKAT-O gene association results comparing patient groups against controls provided below thick line.

| Metric | Controls (n = 166) |  | IBD (n = 282) |  |  | UC (n = 100) |  |  | CD (n = 172) |  |  |
| --- | --- | --- | --- | --- | --- | --- | --- | --- | --- | --- | --- |
|  | max | mean | max | mean | Mann-Whitney U comparison against controls | max | mean | Mann-Whitney U comparison against controls | max | mean | Mann-Whitney U comparison against controls |
| CADD | 2.71 | 0.28 | 3.52 | 0.40 | $1.04 \times 10^{-1}$ | 2.66 | 0.20 | $1.38 \times 10^{-1}$ | 3.52 | 0.54 | $4.62 \times 10^{-4} **$ |
| DANN | 5.92 | 0.84 | 7.62 | 1.06 | $1.36 \times 10^{-1}$ | 5.62 | 0.57 | $1.22 \times 10^{-1}$ | 7.62 | 1.38 | $8.16 \times 10^{-4} **$ |
| FATHMM | 3.33 | 0.49 | 4.34 | 0.66 | $1.04 \times 10^{-1}$ | 3.14 | 0.38 | $1.47 \times 10^{-1}$ | 4.34 | 0.84 | $4.84 \times 10^{-4} **$ |
| fathmm-MKL | 4.53 | 0.37 | 6.24 | 0.55 | $4.54 \times 10^{-2} *$ | 3.78 | 0.25 | $3.15 \times 10^{-1}$ | 6.24 | 0.76 | $1.79 \times 10^{-4} **$ |
| GERP++_RS | 5.30 | 0.64 | 7.00 | 0.87 | $1.26 \times 10^{-1}$ | 4.95 | 0.42 | $1.27 \times 10^{-1}$ | 7.00 | 1.17 | $6.95 \times 10^{-4} **$ |
| M-CAP | 1.87 | 0.12 | 3.39 | 0.22 | $1.58 \times 10^{-2} *$ | 1.73 | 0.08 | $4.62 \times 10^{-1}$ | 3.39 | 0.32 | $1.37 \times 10^{-4} **$ |
| MetaLR | 2.42 | 0.16 | 3.39 | 0.29 | $2.71 \times 10^{-1}$ | 1.81 | 0.10 | $2.34 \times 10^{-2} *$ | 3.39 | 0.42 | $1.63 \times 10^{-3} **$ |
| MetaSVM | 2.67 | 0.30 | 3.61 | 0.43 | $9.88 \times 10^{-2}$ | 2.50 | 0.22 | $1.50 \times 10^{-1}$ | 3.61 | 0.57 | $4.39 \times 10^{-4} **$ |
| MutationTaster | 4.38 | 0.26 | 5.10 | 0.39 | $4.48 \times 10^{-2} *$ | 2.65 | 0.13 | $4.37 \times 10^{-1}$ | 5.10 | 0.57 | $7.47 \times 10^{-4} **$ |
| phastCons | 4.66 | 0.35 | 5.24 | 0.56 | $2.86 \times 10^{-1}$ | 3.54 | 0.24 | $2.70 \times 10^{-2} *$ | 5.24 | 0.77 | $2.16 \times 10^{-3} **$ |
| phyloP | 6.32 | 1.02 | 7.93 | 1.27 | $1.23 \times 10^{-1}$ | 5.92 | 0.75 | $1.38 \times 10^{-1}$ | 7.93 | 1.62 | $7.09 \times 10^{-4} **$ |
| Polyphen2_HDIV | 5.32 | 0.68 | 7.03 | 0.82 | $2.02 \times 10^{-1}$ | 2.30 | 0.33 | $6.22 \times 10^{-2}$ | 7.03 | 1.13 | $1.20 \times 10^{-3} **$ |
| Polyphen2_HVAR | 4.86 | 0.46 | 5.31 | 0.64 | $1.65 \times 10^{-1}$ | 2.07 | 0.21 | $7.22 \times 10^{-2}$ | 5.31 | 0.92 | $7.90 \times 10^{-4} **$ |
| PROVEAN | 4.33 | 0.66 | 5.23 | 0.86 | $1.04 \times 10^{-1}$ | 4.08 | 0.49 | $1.45 \times 10^{-1}$ | 5.23 | 1.10 | $4.84 \times 10^{-4} **$ |
| SIFT | 5.91 | 0.95 | 7.61 | 1.14 | $1.47 \times 10^{-1}$ | 5.43 | 0.64 | $1.16 \times 10^{-1}$ | 7.61 | 1.47 | $9.64 \times 10^{-4} **$ |
| VEST3 | 3.28 | 0.30 | 4.21 | 0.44 | $1.36 \times 10^{-1}$ | 2.24 | 0.17 | $1.13 \times 10^{-1}$ | 4.21 | 0.62 | $7.48 \times 10^{-4} **$ |
| SKAT-O (all variants) | - | - | $5.41 \times 10^{-1}$ | | | $9.76 \times 10^{-2}$ | | | $3.46 \times 10^{-2} *$ | | |
| SKAT-O (MAF<0.05) | - | - | $4.63 \times 10^{-1}$ | | | $1.37 \times 10^{-1}$ | | | $5.02 \times 10^{-2}$ | | |

Interestingly, similar results were observed for the SKAT-O gene test of association when using all variant frequency data as but not when restricting to rare variation (MAF < 0.05). Importantly, the magnitude of the difference between CD patients and control groups was statistically weaker (p = 0.0346) and less robust to correction for multiple testing.

Although not the purpose of this comparison, we confirmed GenePy whole gene comparison provided statistical evidence two orders of magnitude greater than any single variant association result (Supplementary Table 1).

## Discussion

Next-generation sequencing is a disruptive technology set to transform biological assessment. Globally, it is rapidly integrating into the medical sector with numerous countries already funding whole genome sequencing of patient samples for diagnosis and treatment of rare disease and cancer. Multiple metrics have emerged that aim to annotate individual mutations with a view to sensitively implicating causal versus non-causal variation. However, for common complex diseases where the action of an unknown number of multiple variants converge to increase susceptibility, the molecular assessment of mutation profiles is necessarily less binary. Furthermore, in order to bring interpretation from bench to bedside, it is important that methodology provides discriminatory evidence for individual patients and not just evidence of modest genetic effects between large cohorts.

Herein, we describe the implementation of GenePy representing a novel alternative to examine genetic burden that provides a quantitative measure of the combined burden of mutation across each gene for each individual. The scoring system has the freedom to harness the intrinsic properties of any user-defined variant-level deleteriousness metric. By summing across genes, GenePy further integrates biological information on frequency and zygosity and when being used to examine all genes or subsets thereof, can be corrected for gene length.

Implementation of GenePy reveals the high variance necessary to distinguish mutational burden. The logarithmic distribution confers weight to rare pathogenic variants that is additive across a gene and theoretically limited only by the number of variant sites within that gene. The majority of genes return a GenePy score of zero for any one individual but as most coding variation is rare, the likelihood of observing variation in any one gene is positively correlated with cohort size.

We provide proof of principle that the GenePy score has improved sensitivity to detect clinically meaningful gene perturbations. GenePy performance compares favourably against the most commonly applied gene-based association test optimised for small data sets (SKAT-O). Superiority to detect the subtle effects of genes in complex disease is likely attributable to the additional modelling of innate biological features of mutations. Power to determine significant GenePy score differences between patient and control groups was consistent across sixteen different metrics of variant deleteriousness whereby all concordantly reported a similar level of significance despite differing underlying principles. While GenePy scores generated using M-CAP metric returned the most significant difference in CD patients compared to controls, it is likely that no metric will prove optimal in all situations. The GenePy scoring system can easily accommodate new and improved variant deleteriousness metrics that are constantly evolving with more widespread use and interpretation of NGS data.

As with all large-scale data, GenePy scoring is dependent upon data integrity and elimination of systematic bias or technical artefacts. High quality individual DNA samples must be sequenced to sufficient depth to return confident variant calls. For larger scale analyses using multiple samples, parity of capture kits, sequencing platforms and informatic pipelines must be ensured. While these pre-processing quality control steps and generation of the multi-calling VCF file represent the highest computational burden, GenePy score calculation on cleaned vcf files is amenable to batching and computationally trivial.

Many of the currently available deleteriousness scores implemented herein fail to annotate synonymous, splicing or protein truncating variation. While we arbitrarily imposed maximum deleteriousness scores to protein truncating mutations, we standardised the set of variants examined across metrics by excluding synonymous and splicing variants from this analysis. Deleteriousness metrics based on conservation alone are calculable for all genomic variation and could be implemented for the assessment sliding windows of non-coding regions derived from whole-genome sequencing. Due to association testing in Caucasian samples only, we restricted allele frequency annotation to that ethnic group. Arguably, there is merit in the implementation of global allele frequency estimates or those from more ancestrally diverse populations.

Further refinements of the GenePy scoring system might be realised by integration of gene essentiality [49] (and conversely gene redundancy) or gene damage indices (GDI) [50]. Long read NGS data enabling the discrimination of gametic phase would substantially advantage integration of inheritance models and haploinsufficiency.

The key advantage of GenePy is its provision of a continuous quantitative measure of biological integrity of a gene within individuals, resulting in a score that is easily integrated into downstream analyses. GenePy scores are not dependent on cohort size and can be calculated and assessed on a per-patient basis. GenePy scores are suited to pathway analyses where scores can be overlaid and summed across defined molecular cascades. For the particular assessment of complex disease, machine learning tools that integrate multi-omic and extensive biomarker ’big data’ to determine cryptic patterns are increasingly applied. The ability to input biologically rich information and the gene and individual level represents an important step change from the more traditional methods of assessing genetic data at the variant and cohort level.

## Acknowledgments

The authors would like to thank Rachel Haggarty for assistance with management of the genetics of PIBD study database. We also would like to acknowledge Nikki Graham for assistance with sample extraction and management. We thank the EUCLIDS consortium, for providing access to anonymised exome data used for comparison and development of our model.

